# Modulating pathogenesis with Mobile-CRISPRi

**DOI:** 10.1101/618637

**Authors:** Jiuxin Qu, Neha K. Prasad, Michelle A. Yu, Mark R. Looney, Shuyan Chen, Amy Lyden, Emily Crawford, Melanie R. Silvis, Jason M. Peters, Oren S. Rosenberg

## Abstract

Pathogens express a set of proteins required for establishing and maintaining an infection, termed virulence life-style genes (VLGs). Due to their outsized importance in pathogenesis, VLG products are attractive targets for the next generation of antimicrobials. However, precise manipulation of VLG expression in the context of infection is technically challenging, limiting our ability to understand the roles of VLGs in pathogenesis and accordingly design effective inhibitors. We previously developed a suite of gene knockdown tools that are transferred by conjugation and stably integrate into pathogen genomes that we call “Mobile-CRISPRi”. Here we show the efficacy of Mobile-CRISPRi in controlling VLG expression in a murine infection model. We optimize Mobile-CRISPRi in *Pseudomonas aeruginosa* for use in a murine model of pneumonia by tuning the expression of CRISPRi components to avoid non-specific toxicity. As a proof of principle, we demonstrate that knockdown of a VLG encoding the type III secretion system (T3SS) activator ExsA blocks effector protein secretion in culture and attenuates virulence in mice. We anticipate that Mobile-CRISPRi will be a valuable tool to probe the function of VLGs across many bacterial species and pathogenesis models.

**Importance:** Antibiotic resistance is a growing threat to global health. To optimize the use of our existing antibiotics and identify new targets for future inhibitors, understanding the fundamental drivers of bacterial growth in the context of the host immune response is paramount. Historically these genetic drivers have been difficult to manipulate, as they are requisite for pathogen survival. Here, we provide the first application of Mobile-CRISPRi to study virulence life-style genes in mouse models of lung infection through partial gene perturbation. We envision the use of Mobile-CRISPRi in future pathogenesis models and antibiotic target discovery efforts.

## Introduction

All pathogenic bacteria require virulence life-style genes (VLGs) for survival in the host environment (**1**). These include both classical virulence factors that are dispensable for growth in rich media and essential genes that modulate pathogenesis (**2**). Next generation of sequencing of bacterial transposon mutant libraries (e.g. Tn-Seq (**3**), INSeq (**4**)) from infected animals has enabled comprehensive identification of non-essential VLGs in a single experiment, rapidly increasing our knowledge of which genes are required for pathogenesis (**5–16**). There are two major limitations of Tn-Seq to study VLGs, both arising from the complete loss of function usually caused by Tn-mutagenesis. First, core essential genes are by definition excluded from the analysis of environment-specific essentiality. Secondly, all-or-nothing mutations preclude our ability to observe the relationship between expression of the gene product and fitness in the host environment; this information could be valuable in identifying VLGs for which the organism is highly sensitive to slight perturbation, which would be ideal candidates for inhibitors. Thus, methods that can partially perturb VLG function in the context of pathogenesis are highly valuable.

Gene repression tools that are currently used to study VLGs during infection have provided numerous insights into gene function but have key technical limitations. Antisense RNAs (**17, 18**) have variable efficacy, substantial off-target effects (**19–21**), and cannot be rationally designed (**22**). Methods to trigger protein degradation (i.e., degrons (**23–26**)) require each gene of interest to be tagged at its native locus and suffer from toxicity due to interference with protein function and stability (**26**). Gene depletion from inducible promoters also requires insertion of the promoter upstream of all genes of interest and is limited by the inability to optimize both control of non-induced promoter expression (leakiness) and maximal amount of induced gene product (**27**).

In contrast, CRISPRi—use of a catalytically-inactive variant of the Cas9 nuclease (dCas9) to repress transcription (**28**)—is highly efficacious and specific in bacteria (**29**), easily programmable by editing the first 20 nt of the guide RNA (sgRNA (**30**)), does not require modification of the chromosome at each targeted gene, and maintains the native regulation of targeted genes. We previously developed “Mobile-CRISPRi,” a technology that enables transfer and stable integration of CRISPRi systems into diverse bacteria (**Fig. 1A**) (**31**). Here, we optimize Mobile-CRISPRi for targeting VLGs in a *P. aeruginosa* PA14 murine pneumonia model of infection.

**Figure 1.**
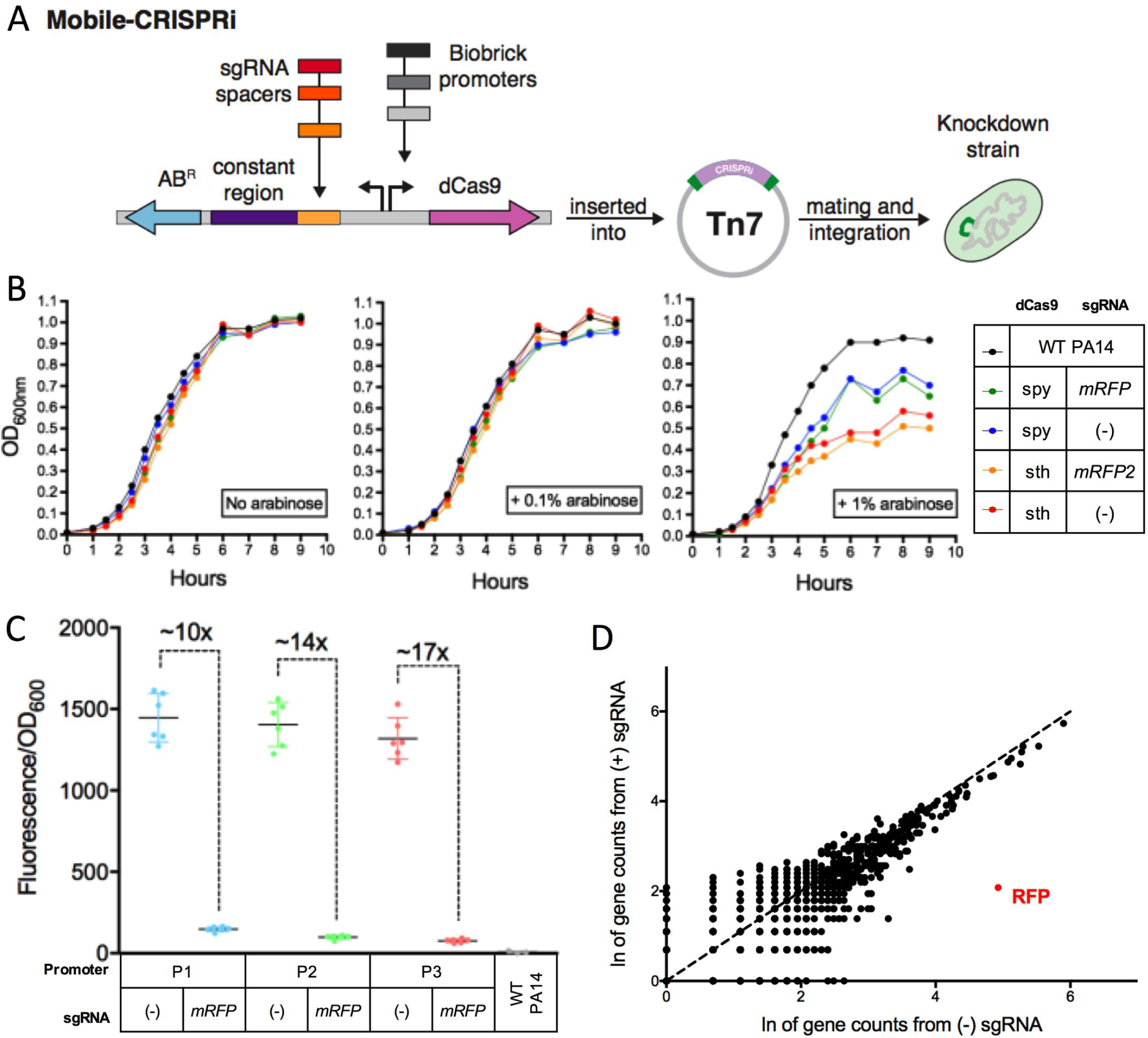
Toxicity, efficacy, and specificity of Mobile-CRISPRi in *Pseudomonas aeruginosa*. **(A)** The pMobile-CRISPRi plasmid is comprised of an antibiotic resistance cassette (AB^R^), sgRNA spacers specific to gene of interest, a promoter driving *dcas9* expression, and *dcas9.* These components can be manipulated before chromosomal integration into a pathogen to generate a hypomorphic strain. **(B)** Wildtype PA14 growth was compared to that of mutant PA14 strains with integrated Mobile-CRISPRi systems featuring arabinose-inducible promoters driving dCas9 activity, two different variants of dCas9 (*S. pyogenes* and *S. thermophilus*), and the presence or absence of *mRFP*-targeting sgRNA. To induce the promoter, these strains were incubated with no arabinose, 0.1 % arabinose, or 1 % arabinose. **(C)** *mRFP* was cloned into Mobile-CRISPRi plasmids under the control of three different constitutive promoters. The median fluorescence of strains without sgRNA was compared to that of strains with mRFP-targeting sgRNA after 14 hours of growth (*p*<.0001 for all three). **(D)** RNA was extracted from mutants featuring P3 with and without mRFP-targeting sgRNA. Gene counts from RNA-seq are plotted for each strain.

### Optimized dCas9 expression eliminates toxicity and allows for graded knockdowns

dCas9 overexpression often causes non-specific toxicity in bacteria (**32**), which would likely complicate the interpretation of our CRISPRi experiments in infection models. Indeed, we found that full induction of an arabinose-inducible promoter (P_*BAD*_) driving the expression of dCas9 variants from *Streptococcus pyogenes* (dCas9_*spy*_) (**28**) or *Streptococcus thermophilus* (dCas9_*sth*_) (**33**) resulted in reduced growth of PA14 in rich culture medium, whereas partial induction showed no apparent toxicity (**Fig. 1B**). We reasoned that titrating chemical inducers (e.g. arabinose) in a mouse model could be impractical due to variable tissue penetration (**34–38**), so we instead focused on expressing dCas9_*spy*_ from a series of weak constitutive promoters from the BioBrick Registry (**39**) to reduce toxicity (**Fig. S1**) and achieve partial knockdown. To assess CRISPRi efficacy using the BioBrick promoter strains, we employed a “test” version of Mobile-CRISPRi expressing monomeric Red Fluorescent Protein (*mRFP*) and an sgRNA targeting the *mRFP* gene (**31**). Knockdown levels were quantified for each promoter through comparing the mutants’ fluorescence normalized to growth over time After 12 hours, we found stable fluorescence ratios between mutants without and with sgRNA **(Fig. S2).** The gradient of knockdown from 10-17-fold at the 14-hour time point roughly corresponds to the BioBrick promoter strength used to express dCas9 (**Fig. 1C**). We performed RNA-seq on cells expressing dCas9 from the strongest of the three BioBrick promoters in our set and confirmed that CRISPRi retained specificity (**Fig. 1D**). We conclude that Mobile-CRISPRi optimized with BioBrick promoters driving dCas9_spy_ enables a non-toxic gradient of constitutive knockdowns in PA14.

### Mobile-CRISPRi targeting of VLGs in a murine pneumonia model

A major goal for developing Mobile-CRISPRi in infection models is to identify VLGs for which a modest perturbation has a substantial impact on pathogenesis. To do so, the system must demonstrate stable repression of the gene of interest over the course of infection. As a test case, we targeted *exsA*, which encodes the key activator of T3SS genes in *P. aeruginosa*. Because the *exsA* gene is positively autoregulated (**40**), we reasoned modest knockdown would cause a large reduction in transcription of T3SS genes, resulting in a loss of effector secretion and impaired virulence. Consistent with this, we found that CRISPRi knockdown of *exsA* reduced expression of T3SS genes by more than 100-fold (**Fig. 2A**), similar to expression levels observed in a strain with an *exsA* disruption (*exsA*::Tn (**41**)). As anticipated, knockdown of a positively autoregulated transcriptional factor suppressed the previously observed promoter-dependent gradation. Of note, the high levels of *exsA* in the *exsA*::Tn strain is attributed to amplifying the region of the gene upstream of transposon insertion (**Table S3**). Knockdown of *exsA* also eliminated detectable production of T3SS pilus (PopB/D) and effector (ExoT/U) proteins (**Fig. 2B**). Neither the *exsA* knockdown nor the non-targeting control sgRNA strains showed a growth defect in rich culture medium (**Fig. S1**).

**Figure 2:**
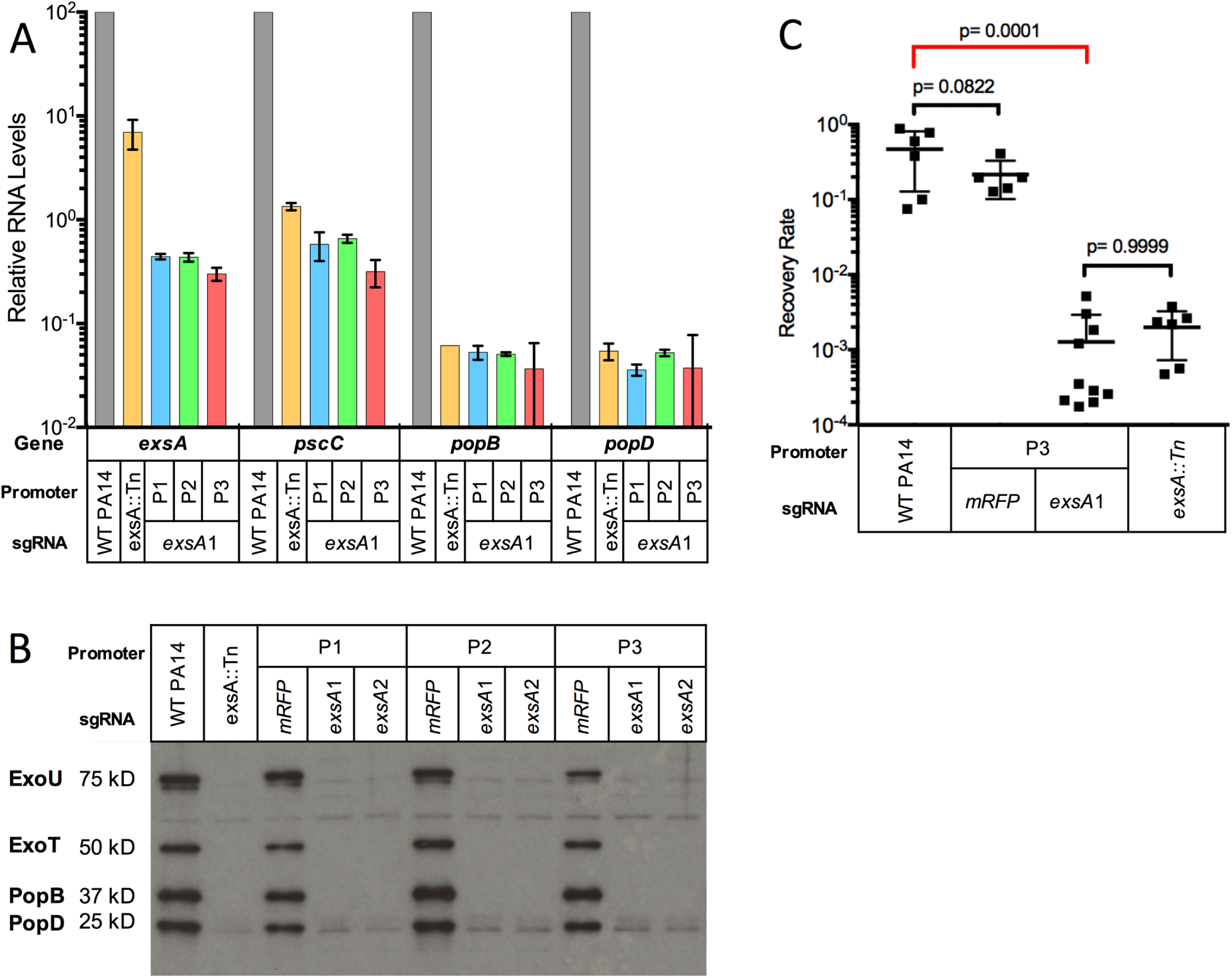
T3SS-associated gene transcription, protein secretion profiles, and clearance of PA14 strains in infection model. **(A)** QRT-PCR analysis for T3SS-related genes across PA14 strains, normalized to WT PA14 RNA levels. **(B)** Immunoblot analysis of exoenzyme U and exoenzyme S secretion for PA14 strains grown in MinS medium (for type III protein secretion induction). **(C)** Following infection, lung homogenate serial dilutions were plated to estimate the CFU of bacteria recovered from the lung. Recovery rate is output CFU relative to input CFU.

Loss of *exsA* function is known to strongly attenuate virulence in a murine pneumonia model (**42, 43**). To test whether Mobile-CRISPRi can be used to probe the functions of VLGs such as *exsA* in a host environment, we intratracheally instilled C57BL/6 mice with a range of 10^5^ to 10^7^ CFU of WT PA14, an isogenic *exsA*::Tn mutant, or Mobile-CRISPRi mutants containing either an sgRNA targeting *exsA* or a non-targeting control. We collected the lungs 18 hours after infection and plated lung homogenates to estimate the number of viable bacteria (**Fig. S2**) (**44**). Strains with the *exsA*::Tn allele or Mobile-CRISPRi targeting *exsA* were highly attenuated for virulence and yielded similar recovery rates. This demonstrates that Mobile-CRISPRi is an effective tool to knockdown VLGs in PA14 during a mouse infection and implies that Mobile-CRISPRi is as stable during *in vivo* infection as it is during growth in culture (**31**). Furthermore, CFU recovery was similar between WT and non-targeting Mobile-CRISPRi, suggesting that non-specific toxicity of dCas9 was mitigated by reduced expression. Other general indicators of infection including hypothermia and leukopenia were observed for the non-targeting and WT controls. These phenotypes were similarly diminished among the *exsA* targeting strain, *exsA* disruption strain, and PBS control (**Fig. S3**). We conclude that Mobile-CRISPRi can probe VLG phenotypes in infection models.

## Discussion

We previously demonstrated that Mobile-CRISPRi could be used to repress gene expression in a number of bacterial pathogens associated with antibiotic resistance (e.g., the ESKAPE pathogens: *Enterococcus, Staphylococcus, Klebsiella, Acinetobacter, Pseudomonas*, and *Enterobacter* (**31, 45**)). Our optimized Mobile-CRISPRi system opens the door to systematic analysis of VLGs in these pathogens during infection, enabling drug-gene interaction studies and, in principle, a screen for new inhibitors that synergize with the host immune system.

## Methods

### Construction of Mobile-CRISPRi plasmids and strains

Plasmids encoding nuclease-null *Streptococcus pyogenes* and *Streptococcus thermophilus* dCas9s were gifted by Lei Qi and Sarah Fortune, respectively. The vectors containing Tn7-based Mobile-CRISPRi system were constructed as previously described by Peters (**31**). dCas9 was expressed from arabinose-inducible P_BAD_ promoter and three constitutive promoters: Anderson BBa_J23117 (P1), Anderson BBa_J23114 (P2) and Anderson BBa_J23115 (P3). The chimeric sgRNA was expressed by a constitutive promoter, P_trc_ with no LacI operator site. In this study, all the mutants were constructed from *Pseudomonas aeruginosa* UCBPP-PA14 by tri-parental mating as previously described (**31**). A complete list of plasmids and strains used in the study can be found in **Supplementary Table 1 and 2**, respectively. PA14 *exsA*::Tn strain was obtained from a transposon insertion library (**41**).

### Toxicity measurements

For dCas9 toxicity measurements, WT PA14 and the mutants were streaked on Pseudomonas Isolation Agar (PIA) plates and incubated for 20 hours at 37°C. On the second day, 2 ml LB medium tube containing one colony from each plate were cultured in a shaker incubator at 37°C and 350 rpm for 12 hours. Then cultures were diluted in 100 μL LB medium with no inducer, 0.1% arabinose, or 1% arabinose to yield a mixture with OD_600nm_ of 0.05 in a 96-well plate (Cat. No. 351177, Corning, NY). These were grown with a lid for 9 - 10 hours on a plate shaker (OrbiShaker MP, Benchmark Scientific, NJ) at 37°C and 900 rpm. The OD_600nm_ of the plate cultures were measured every hour using a microplate reader (SpectraMax 340PC, Molecular Devices, CA).

### RFP knockdown efficiency

Following tri-parental mating, two *P. aeruginosa* colonies were picked from each strain to serve as biological replicates and were incubated overnight in 3 mL of LB + 100 μg/mL gentamicin selective media at 37°C with shaking. These cultures were diluted to .01 OD_600_ into fresh LB media, and 200 μL of this culture was added in triplicate to a clear bottom, black 96 well plate (Corning Costar®). This was covered with an optically clear seal, and a needle was used to poke holes in each of the wells. Fluorescence (excitation: 557nm, emission: 592nm) and OD_600_ were monitored during incubation in a microplate reader (Synergy H1, BioTek Instruments, VT) with continuous, fast, double orbital shaking. Samples were blanked with a well containing LB media. For each replicate, fluorescence value was divided by OD_600_ values at each time-point and plotted in 30-minute intervals. Ratios of median fluorescence from strains ((-):(+) sgRNA per promoter) were averaged between 12 and 18 hours to calculate knockdown ratios.

### RNA extraction

PA14 Mobile-CRISPRi mutants, *exsA*::Tn7, and WT were streaked onto VBMM or LB agar plates and incubated overnight at 37°C. One colony from each plate was grown in MinS (T3SS-inducing minimal media supplemented with nitrotriacetic acid and lacking calcium medium) (**46**) or LB medium at 37°C for 16 hours with shaking at 250 rpm. Then the strains were subcultured in 400 µL fresh MinS or LB medium until the OD_600nm_ reached 1.0. Total RNA was extracted from cell pellets using the RNeasy minikit (Qiagen) according to the manufacturer’s instructions with on-column DNase I digestion (Qiagen). The RNA extracts were aliquoted and stored at - 80°C.

### Quantitative reverse transcriptase PCR (qRT-PCR)

cDNA was synthesized using Random Hexamer primers and RevertAid First Strand cDNA Synthesis Kit (Thermo Scientific; Waltham, MA). To check the amplification efficiency of the primers, 1:50 dilution of WT PA14 cDNA was mixed with “PowerUp SYBR Green Master Mix” (Thermo Scientific; Waltham, MA) and detected by the MX3000P qPCR System (Stratagene, La Jolla, CA, USA). Primers for T3SS related genes (**Table S3)** had amplification efficiency between 90%-110%. PA14 housekeeping gene, *nadB*, was used as internal control for normalization of total RNA levels (**47**). The relative efficiency of each primer pair was tested and compared with that of *nadB* and the threshold cycle data analysis (2^-ΔΔCt^) was used (**48**). All reactions were performed in triplicates and repeated at least twice using independent cultures, with average values of biological replicates and error bars representing standard deviation of ΔΔCt.

### cDNA library preparation and RNA-Seq

The RNA concentration for each sample was determined with a NanoDrop spectrophotometer (Thermo Fisher Scientific). 10 ng total RNA of each sample was fragmented for 6 min, cDNA libraries were prepared using “NEBNext^®^ Ultra™ RNA Library Prep Kit for Illumina^®^” (NEB#E7770S®). Libraries were sequenced in collaboration with the Chan Zuckerberg Biohub in San Francisco on an Illumina MiSeq in 150 bp paired-end runs. Approximately 1,000,000 reads were collected for each of the two samples, with ∼94% alignment to PA14 WT by Bowtie2 (**49**), and transcripts were counted with HTSeq (**50**).

### Type III Secretion Profile of *Pseudomonas aeruginosa* by Immunoblotting

To knockdown *exsA* gene, two specific sgRNAs, *exsA*1 and *exsA*2 were designed. PA14 Mobile-CRISPRi mutants, *exsA::*Tn, and WT were streaked onto VBMM agar plates and incubated at 37°C for overnight. One colony from each plate was grown at 37°C for 16 hours in a shaking incubator at 250 rpm in MinS media (**46**). Bacteria were removed by centrifugation at 6000 × g for 15 min. Then the supernatant was collected and the secreted proteins were precipitated by the addition of ammonium sulfate. The protein pellets were dissolved in sample buffer. After boiling, samples were loaded onto ExpressPlus 4-20% PAGE gels (Genscript; Piscataway, NJ) and run under denaturing conditions. PAGE gels were transferred to PVDF membrane and immunoblotted with polyclonal rabbit antiserum against ExoU, ExoT/ExoS, PopB and PopD proteins as previously described (**46**).

### Murine Infection Model

Pathogen-free male C57BL/6J mice, 8 weeks of age, were purchased from Jackson Laboratories. Animal experiments were conducted in accordance with the approval of the Institutional Animal Care and Use Committee (IACUC) at UCSF. 29 mice were randomly assigned in five groups, G1: WT PA14, 6 mice; G2: P3 with sgRNA *mRFP*, 5 mice; G3: P3 with *exsA*1 sgRNA, 10 mice; G4: *exsA*::Tn, 6 mice; G5: saline control, 2 mice. Mice were anesthetized with isofluorane prior to intratracheal instillation with bacteria at a range of 1×10^5^ to 1×10^7^ CFU/animal in a volume of 50 mL per an established protocol (**44**). Animal weights and rectal temperatures were measured prior to euthanasia. The lungs were collected in 1mL of sterile PBS and processed with a handheld homogenizer (Kinematica, Polytron PT1200E). 50 μl of lung homogenate with appropriate dilutions were spread onto PIA plates with and without gentamicin to count output CFU. Bacterial recovery rate was calculated as the ratio of output CFU to input CFU. For whole blood analysis, blood was collected by cardiac puncture into acid citrate dextrose (Sigma-Aldrich), and WBC was measured by hematology analyzer (Genesis, Oxford Science).

### Statistical analysis

GraphPad Prism (v. 7.0) was used for the statistical analysis of all the data. *P* values < 0.05 were considered statistically significant. For the RFP fluorescence assay, Ordinary one-way ANOVA followed by Turkey’s multiple comparison test was used to compare OD_600_-normalized median fluorescence values between strains featuring the same promoter with and without sgRNA. Ordinary one-way ANOVA followed by Turkey’s multiple comparison test was also used to compare log-transformed bacterial recovery rate, temperature, WBC counts, and weight changes.

## Acknowledgements

We thank S. Fortune (Harvard University) for dCas9_sth_ plasmid, A. Hauser (Northwestern University) for polyclonal rabbit antiserum for *exsA*-related proteins, and C. Gross (UCSF) and J. Engel (UCSF) for productive discussions and some strains. The sequencing team at the Chan Zuckerberg Biohub (N. Neff, B. Yu, R. Sit, and M. Tan) assisted with RNAseq experiment and analysis. C. Mahendra and A. Borges in the lab of J. Bondy-Demony (UCSF) provided technical assistance during the fluorescence plate reader assay. This work was supported by NIH 1K22AI137122 and Innovative Genomics Institute (to J.M.P.), NIH R35 GM118061 (to Carol A Gross to support M.S.), NIAID R01 AI125445 (to M.R.L.), NSF-GRFP 1650113 (to N.K.P.), the National Natural Science Foundation of China 81400005/H0104 and Special Support Fund of Shenzhen for Introduced High-Level Medical Team and Translational medicine of Biochip in clinical laboratory SZSM201412005 (to J.Q.), and 1R01AI128214, 5R01EB024014, P01AI095208, U19AI135990, Chan-Zuckerberg Biohub, CF Foundation Research Development Program, and Gilead Sciences Research Scholars Program in Cystic Fibrosis (to O.S.R).

